# Heritability Informed Power Optimization (HIPO) Leads to Enhanced Detection of Genetic Associations Across Multiple Traits

**DOI:** 10.1101/218404

**Authors:** Guanghao Qi, Nilanjan Chatterjee

## Abstract

Genome-wide association studies have shown that pleiotropy is a common phenomenon that can potentially be exploited for enhanced detection of susceptibility loci. We propose heritability informed power optimization (HIPO) for conducting powerful pleiotropic analysis using summary-level association statistics. We find optimal linear combinations of association coefficients across traits that are expected to maximize non-centrality parameter for the underlying test statistics, taking into account estimates of heritability, sample size variations and overlaps across the traits. Simulation studies show that the proposed method has correct type I error, robust to population stratification and leads to desired genome-wide enrichment of association signals. Application of the proposed method to publicly available data for three groups of genetically related traits, *lipids (N=188,577)*, *psychiatric diseases (N*_*case*_*=33,332, N*_*control*_*=27,888)* and *social science traits* (*N* ranging between 161,460 to 298,420 across individual traits) increased the number of genome-wide significant loci by 12%, 200% and 50%, respectively, compared to those found by analysis of individual traits. Evidence of replication is present for many of these loci in subsequent larger studies for individual traits. HIPO can potentially be extended to high-dimensional phenotypes as a way of dimension reduction to maximize power for subsequent genetic association testing.

## Introduction

Genome-wide association studies of increasingly large sample sizes are continuing to inform genetic basis of complex diseases. These studies have now led to identification of scores of susceptibility SNPs underlying a vast variety of individual complex traits and diseases^1–3^. Moreover, analyses of heritability and effect-size distributions have shown that each trait is likely to be associated with thousands to tens of thousands of additional susceptibility variants, each of which individually has very small effects, but in combinations they can explain substantial fraction of trait variation^4–15^. GWAS of increasing sample sizes as well as re-analysis of current studies with powerful statistical methods are expected to lead to identification of many of these additional variants.

An approach to increase the power of existing GWAS is to borrow strength across related traits. Comparisons of GWAS discoveries across traits have clearly shown that pleiotropy is a common phenomenon^3,14,16–19^. Aggregated analysis of multiple related traits have led to identification of novel SNPs that could not be detected through analysis of individual traits alone^20–23^. Further, analysis of genetic correlation using genome-wide panel of SNPs have identified groups of traits that are likely to share many underlying genetic variants of small effects^10,12,14,24,25^. As summary-level association statistics from large GWAS are now increasingly accessible, there is a great opportunity to accelerate discoveries through novel cross-trait analysis of these datasets.

A variety of methods have been developed in the past decade to increase power of GWAS analysis by combining information across multiple traits^13,26–37^. Many of these methods have focused on developing test-statistics that are likely to have optimal power for detecting an individual SNP under certain types of alternatives of its shared effects across multiple traits^26,30,31,35,38,39^. These approaches do not borrow information across SNPs and may be inefficient for analysis of traits that are likely to have major overlap in their underlying genetic architecture. For the analysis of psychiatric diseases, for example, it has been shown that borrowing pleiotropic information across SNPs can be used to improve power of detection of individual SNP associations and genetic risk prediction^40,41^.

In this article, we propose a novel method for powerful aggregated association analysis for individual SNPs across groups of multiple, highly related, traits informed by genome-wide estimates of genetic variance-covariance matrices. We derive optimal test-statistics based on orthogonal linear combinations of association coefficients across traits that are expected to maximize genome-wide averages of the underlying non-centrality parameters in a gradually decreasing order. We exploit recent developments in LD-score regression methodology^14,42^ for estimation of phenotypic and genotypic correlations for implementation of the method using only summary-level results from GWAS.

We evaluate performance of the proposed method through extensive simulation studies using a novel scheme for directly generating summary-level association statistics for large GWAS for multiple traits with possibly overlapping samples. We use the proposed method to analyze summary-statistics available from consortia of GWAS of lipid traits^43^, psychiatric diseases^20^ and social science traits^44^. These applications empirically illustrate that HIPO directions can be highly enriched with association signals and can identify novel and replicable associations that are not identifiable at comparable level of significance based on analysis of the individual traits.

## Material and Methods

### Model and Assumptions

Suppose that the summary level results are available for *K* traits. For a given SNP *j*, let 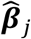 and *s*_*j*_ denote vectors of length *K* containing estimates of regression parameters and associated standard errors, respectively, for the *K* traits. Let *M* be the total number of SNPs under study. Throughout, we will assume both genotypes and phenotypes are standardized to have mean 0 and variance 1. Let *N*_k_ denote the sample size for GWAS for the *k*-th trait. We assume *N*_k_ can vary across studies because traits may be measured on distinct, but potentially overlapping, samples. We assume that summary-level statistics in GWAS are obtained based on one SNP at a time analysis and that 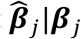 follows a multivariate normal distribution: 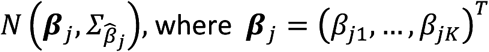 is referred to as the “marginal” effect sizes, the coefficients that will be obtained by fitting single-SNP regression models across the individual traits in the underlying population.

The variance-covariance matric 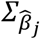 which may include non-zero covariance terms when the studies have overlapping samples, will be estimated based on estimates of standard errors of the individual coefficients (*s*_*j*_) and estimate of “phenotypic correlation” that could be obtained based on LD-score regression.

### Power Optimization

Power has a one-to-one correspondence with the non-centrality parameter (NCP, denoted by *δ*) of the underlying *χ*^2^-statistic. Therefore, we try to find the linear combination 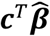 that maximizes the average NCP across SNPs (denoted by *E*[*δ*]), which is given by

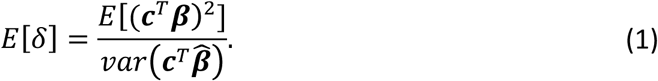

The denominator is easy to simplify: 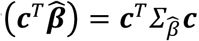 which does not depend on true value of ***β***. We derive an expression of the numerator based on commonly used random effect models that are used to characterize genetic variance-covariances.

Let 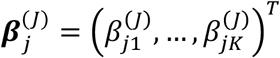 denote the vector of “joint” effect sizes associated with SNP *j* that could be obtained by simultaneous analysis of SNPs in multivariate models across the K individual traits. We assume that 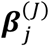 follows a multivariate normal distribution *N* 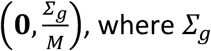 is the genetic covariance matrix. It follows that ***β*_*j*_**, the vector of marginal regression coefficients, is also normally distributed with mean **0** and 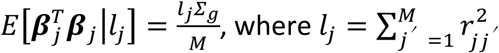 is the LD score. Here 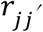 is the correlation of genotypes between SNP 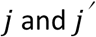.

Thus, based on the above model, the numerator of (1) can be written as 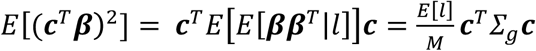. Therefore, we have

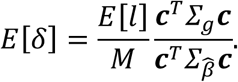

The matrix 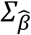 needs to take into account the sample size differences and overlaps across studies. When all the phenotypes are measured on the same set of people, 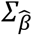 is proportional to the phenotypic variance-covariance matrix and *E[δ]* reduces to maximizing the heritability (MaxH)^34^. But HIPO is more general and can be applied to traits measured on different samples with unknown overlap. The LD-score regression allows estimation of both 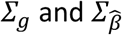 based on underlying slope and intercept parameters, respectively, using GWAS summary-level statistics (**Appendix A**)^14,42^.

The first HIPO component ***c***_*1*_ is given by solving the following optimization problem:

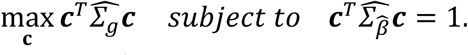

Subsequent components *c*_*k*_ are defined iteratively by solving a slightly different optimization problem

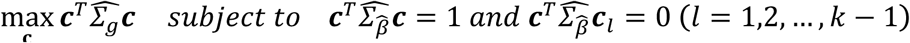

The above procedure can be implemented by suitable eigen decomposition (**Appendix B**). We call the first HIPO component HIPO-D1, the second HIPO component HIPO-D2, and so on. Interestingly, it can be shown that the eigenvalues resulted from this procedure are the average NCP for *χ*^2^ association-statistics across SNPs along the HIPO directions (up to the same scale constant, **Appendix B**). For the *k*th HIPO component, the association for the SNP *j* is tested using *Z*-statistics in the form

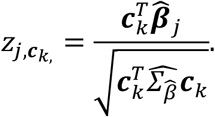

It is easy to see that HIPO *z*-statistics reduce to the inverse standard error weighted *z*-scores when all traits have the same heritability, have genetic correlation 1 and, there is no sample overlap across studies. Therefore, HIPO can also be viewed as an extension of standard meta-analysis.

### Simulations

We use a novel simulation method that directly generates summary level data for GWAS of multiple traits preserving realistic genotypic and phenotypic correlation structures.

We proposed the single-trait version of this approach in a recent study^15^. We propose to simulate GWAS estimate for marginal effects across *K* traits, denoted as 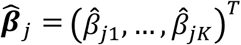 using a model of the form 

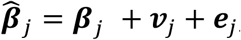

where two types of errors terms, ***ν***_*j*_ and ***e***_*j*_, are introduced to account for variability due to population stratification effects and estimation uncertainty, respectively. We assume the population stratification effects ***ν***_*j*_s follow i.i.d. multivariate normal distribution across SNPs. We generate the estimation error terms 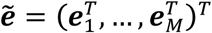 following a multivariate normal distribution that takes into account both phenotypic correlation across traits and linkage disequilibrium across SNPs. In particular we generate 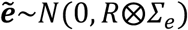 where the covariance matrix is the Kronecker product of the LD
coefficient matrix 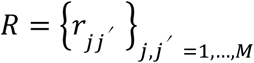 and

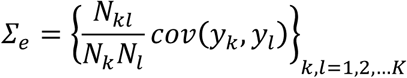

where the *(k, l)* element involves sample sizes, the sample overlap *N*_*kI*_ and the phenotypic covariance between the *k*th and *l*th trait (**Appendix C**). We assume that the sample size is the same for all the SNPs within the same study.

We simulate ***β***_*j*_ by first randomly selecting ~12K causal SNPs out of a reference panel of ~1.2 million HapMap3 SNPs lwith MAF >5% in 1000 Genomes European population. This SNP list is downloaded from LD Hub^45^. For selected casual SNPs, we generate i.i.d. joint effect sizes 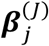 from a multivariate normal distribution 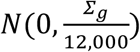 where *∑*_*g*_ is the genetic covariance matrix. For simplicity we assume all the traits have the same set of causal SNPs. We calculate the marginal effect sizes ***β***_*j*_ as the sum of the joint effect size of SNPs in neighborhood 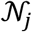 weighted by the LD coefficient, i.e. 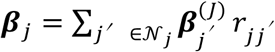. The neighborhood 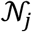 is defined to be set of SNPs that are within 1MB distance and have *r*^2^ > 0.01 with respect to SNP *j*.

For simulation of 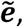 we observe that in a GWAS study where the phenotypes have no association with any of the markers, the summary-level association statistics is expected to follow the same multivariate distribution as 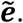 We utilize individual level genotype data available from 489 European samples from the 1000 Genomes Project. For each of the 489 subjects, we simulate a vector of phenotype from a predetermined multivariate normal distribution without any reference to their genotypes. We then conducted standard one SNP at a time GWAS analysis for each trait to compute the association statistics 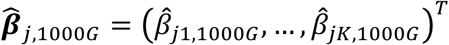 for the 1.2 million SNPs. To mimic the incomplete sample overlap between traits, we can calculate 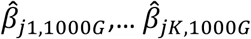 based on different subsamples of 1000 Genomes EUR, of size *n*_1_, …, *n*_k_. Finally, to generate error terms according to sample size specification for our simulation studies, we use the adjustment

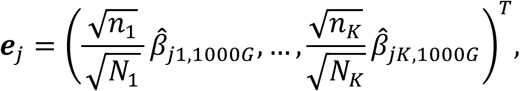

We show in **Appendix C** that this 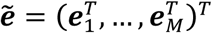 has the desired distribution.

We conduct simulation studies to validate HIPO-based association tests and investigate expected power gain under varying sample size and heritability. For simplicity, we only consider the scenarios where all traits are measured on the same set of subjects. To make the settings more realistic, we use two sets of genetic and phenotypic covariance matrices estimated from real data:

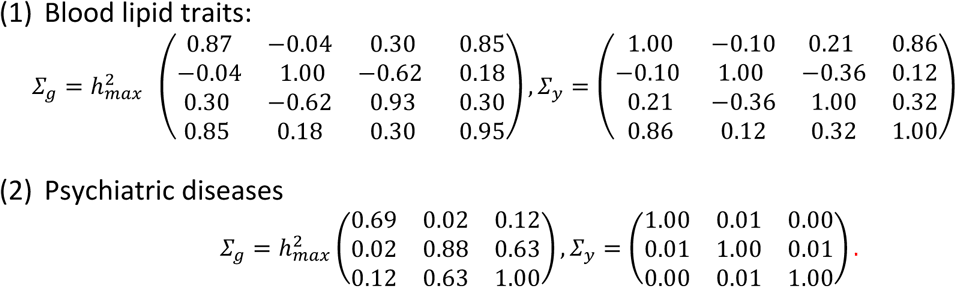

We vary the value of scale factor 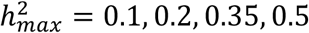 to control heritability of the traits while preserving the genetic correlation structure. We also vary the sample size: *N* = 10K, 50K, 100K, 500K. The covariance matrix of *ν*_*j*_ is set to

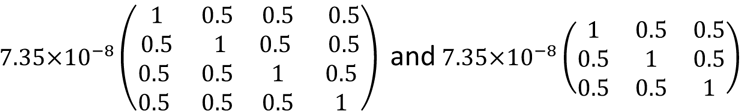

in the first and second settings, respectively. This choice of parameters lead to an average per-SNP population stratification that is about 25% of the per SNP heritability when 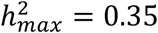 For each of the 2*4*4 = 32 settings we repeat the simulation 100 times.

### Summary level data

We analyze publicly available GWAS summary-level results across three groups of traits measured on European ancestry samples using the proposed method. Global Lipids Genetics Consortium (GLGC) provides the GWAS results for levels of low-density lipoprotein (LDL) cholesterol, high-density lipoprotein (HDL) cholesterol, triglycerides (TG) and total cholesterol (TC)^43^. The data consists of 188,577 European-ancestry individuals with ~1.8 million SNPs after implementing the LD Hub quality control procedure (described at the end of this section).

The Psychiatric Genomics Consortium (PGC) cross-disorder study analyzed data for 5 psychiatric disorders: autism spectrum disorder (ASD), attention deficit-hyperactivity disorder (ADHD), bipolar disorder (BIP), major depressive disorder (MDD) and schizophrenia (SCZ)^20,46–49^. Two of the five traits involved trio data: ASD (4788 trio cases, 4788 trio pseudocontrols, 161 cases, 526 controls, equivalent to 4949 cases and 5314 controls) and ADHD (1947 trio cases, 1947 trio pseudocontrols, 840 cases, 688 controls, equivalent to 2787 cases and 2635 controls). The other three studies did not involve trios: BIP (6990 cases, 4820 controls), MDD (9227 cases, 7383 controls) and SCZ (9379 cases, 7736 controls). After applying the same QC procedure, we included ~1.05 million SNPs for HIPO analysis.

The Social Science Genetic Association Consortium (SSGAC) provides summary statistics for depressive symptoms (DS, N=161,460), neuroticism (NEU, N=170,911) and subjective well-being (SWB, N=298,420)^44^. The DS data is the meta-analysis result combining a study by the Psychiatric Genomics Consortium^48^, the initial release of UK Biobank (UKB)^50^ and the Resource for Genetic Epidemiology Research on Aging cohort (dbGap, phs000674.v1.p1). For neuroticism, the study pooled summary level data sets from UKB and Genetics Personality Consortium (GPC). The SWB data is the meta-analysis result from 59 cohorts^44^. All subjects are of European ancestry. We analyzed ~2.1 million SNPs after QC.

For all three groups of traits, we use the GWAS parameter estimates and standard errors to compute the z-statistics and p-values without making post-meta-analysis correction of genomic control factors. We perform SNP filtering to all three groups of phenotypes based on LD Hub quality control guideline. Markers that meet the following conditions are removed: (1) with extremely large effect size (*χ^2^ > 80*) (2) within the major histocompatibility complex (MHC) region (26Mb~34Mb on chromosome 6) (3) MAF less than 5% in 1000 Genomes Project Phase 3 European samples (4) sample size less than 0.67 times the 90th percentile of the sample size (5) alleles do not match the 1000 Genomes alleles. We further remove SNPs that are missing for at least one trait. The summary statistics are supplied to LDSC software^14,42,45^ to fit LD score regression.

We defined a locus to be “novel” if it contains at least one SNP that reach genome-wide significance (p-value < 5×10^-8^ by the HIPO method and the lead SNP in the region is at least 0.5 Mb away and has *r*^2^ < 0.1 from all lead SNPs of genome-wide significance regions identified by individual trait analysis.

## Results

Simulation results show that HIPO-D1 maintains the correct type I error rate with or without population stratification, consistently across different sample sizes and values of heritability (**Supplementary Tables 1 and 2**). In the presence of population stratification, the degree of which was modest according to our simulation scheme, tests based on individual traits showed somewhat inflated type I error under large sample size (e.g. 500K) (**Supplementary Tables 1 and 2**). Results also show that association analysis based on HIPO-D1 leads to substantial number of additional true discoveries compared to that based on the most heritable trait (**Supplementary Table 3**). In most settings, the value of *λ*_*GC*_ and average *χ*^2^-statistics are larger for HIPO-D1 than those for the most heritable trait (**Supplementary Table 4**). Results also show that QQ plots for HIPO-D1 to be more enriched with signals than those for the most heritable traits (**Supplementary Figures 1-4**).

### Application to blood lipid data

We applied our method to the Global Lipids Genetics Consortium (GLGC) data^43^. The average NCP decreases from 0.213 for HIPO-D1 to 0.026 for HIPO-D4, with most association signals appears to be associated with the first and second components (**Supplementary Table 5**). HIPO-D1 is positively related to TG, negatively related to HDL and TC and depends weakly on LDL. HIPO-D2 depends mostly on TC. The last component, which contains very little genetic association signals, is positively correlated with TC and negatively with the other three traits. The order of *λ*_*GC*_ and average of empirical *χ*^2^ statistic also tracks with the average NCP, suggesting that the observed enrichments are likely due to polygenic effects. We identified twenty novel loci by HIPO-D1 and 4 by HIPO-D2 at genome-wide significant level *(p* < 5×10^-8^) (**Table 1**). The pattern of p-values for individual traits show that the proposed method detects novel SNPs that contain moderate degree of association signals across multiple traits. There is very little overlap between new loci found by HIPO-D1 and by HIPO-D2, as expected from genetic orthogonality of the two components (**Supplementary Figure 5**).

**Table 1.** Novel loci discovered at genome-wide significance level (*p* < 5×10^-8^) by the first and second HIPO components of blood lipid traits. Independent SNPs were identified through LD-pruning with *r*^2^ threshold of 0.1 and pruned SNPs were assumed to represent independent loci if they are >0.5Mb apart. Loci are considered novel if they are not identified at genome-wide significance level through analysis of individual traits. For each lead SNP, p-values for association are shown for HIPO components and for individual traits. The directions of association (+/-) are also shown for each of the individual traits.

| SNP | CHR | MBP | Nearest Gene (Distance) | $P_{LDL}$ | $P_{HDL}$ | $P_{TG}$ | $P_{TC}$ | $P_{HIPO-D1}$ | $P_{HIPO-D2}$ |
| --- | --- | --- | --- | --- | --- | --- | --- | --- | --- |
| **HIPO-D1** |  |  |  |  |  |  |  |  |  |
| rs4850047 | 2 | 3.6 | RPS7(+6.244kb) | 2.87e-03 (+) | 1.15e-04 (+) | 8.58e-04 (-) | 2.14e-06 (+) | 1.13e-09 | 7.82e-04 |
| rs2249105 | 2 | 65.3 | CEP68(0) | 8.72e-02 (+) | 6.35e-06 (+) | 1.89e-06 (-) | 4.66e-01 (+) | 1.33e-08 | 7.92e-01 |
| rs2062432 | 3 | 123.1 | ADCY5(0) | 9.78e-01 (+) | 3.88e-06 (-) | 2.44e-03 (+) | 6.75e-02 (-) | 4.30e-08 | 5.52e-01 |
| rs6855363 | 4 | 157.7 | PDGFC(-12.22kb) | 6.44e-01 (-) | 3.20e-07 (+) | 3.18e-04 (-) | 5.89e-01 (+) | 2.27e-08 | 5.44e-01 |
| rs17199964 | 4 | 102.7 | BANK1(-3.972kb) | 3.11e-01 (-) | 9.19e-08 (-) | 1.27e-01 (+) | 9.85e-04 (-) | 3.03e-08 | 2.43e-02 |
| rs10054063 | 5 | 173.4 | CPEB4(+5.085kb) | 7.00e-01 (-) | 6.13e-04 (-) | 7.72e-07 (+) | 2.67e-01 (-) | 3.69e-08 | 7.80e-01 |
| rs11987974 | 8 | 36.8 | KCNU1(+30.17kb) | 7.66e-01 (+) | 3.79e-06 (-) | 1.85e-06 (+) | 7.16e-01 (-) | 3.91e-09 | 3.32e-01 |
| rs740746 | 10 | 115.8 | ADRB1(-11.02kb) | 4.32e-01 (-) | 1.09e-06 (-) | 1.61e-03 (+) | 1.07e-01 (-) | 4.21e-08 | 5.74e-01 |
| rs10832027 | 11 | 13.4 | ARNTL(0) | 2.53e-02 (+) | 1.52e-07 (+) | 5.73e-07 (-) | 1.87e-02 (+) | 3.88e-12 | 3.16e-01 |
| rs7938117 | 11 | 68.6 | CPT1A(0) | 3.69e-01 (-) | 2.05e-07 (-) | 9.49e-06 (+) | 4.01e-02 (-) | 3.35e-11 | 5.30e-01 |
| rs1565228 | 11 | 27.6 | BDNF-AS(0) | 2.46e-01 (-) | 3.50e-05 (-) | 3.44e-06 (+) | 7.19e-02 (-) | 2.56e-09 | 6.44e-01 |
| rs661171 | 11 | 110.0 | ZC3H12C(0) | 9.06e-03 (+) | 9.84e-07 (+) | 1.77e-01 (-) | 2.70e-06 (+) | 2.58e-08 | 2.23e-04 |
| rs895953 | 12 | 122.2 | SETD1B(0) | 7.23e-01 (-) | 1.45e-06 (+) | 3.98e-07 (-) | 5.43e-01 (+) | 2.84e-10 | 3.86e-01 |
| rs2384034 | 12 | 113.2 | RPH3A(-24.86kb) | 9.47e-03 (+) | 3.61e-07 (+) | 3.18e-02 (-) | 5.00e-05 (+) | 4.39e-09 | 2.51e-03 |
| rs11048456 | 12 | 26.5 | ITPR2(-25.2kb) | 8.65e-02 (-) | 2.93e-07 (-) | 1.59e-03 (+) | 5.74e-02 (-) | 1.75e-08 | 3.28e-01 |
| rs721772 | 15 | 41.8 | RPAP1(0) | 5.17e-01 (+) | 2.26e-07 (-) | 4.30e-05 (+) | 7.10e-01 (-) | 4.25e-09 | 3.75e-01 |
| rs11079810 | 17 | 46.2 | SKAP1(0) | 1.99e-02 (+) | 1.71e-07 (+) | 2.23e-04 (-) | 9.51e-03 (+) | 4.30e-10 | 1.32e-01 |
| rs4805755 | 19 | 32.9 | ZNF507(0) | 9.34e-01 (-) | 5.58e-08 (+) | 4.83e-03 (-) | 9.85e-02 (+) | 7.71e-09 | 6.28e-01 |
| rs10408163 | 19 | 47.6 | ZC3H4(0) | 1.47e-01 (-) | 9.99e-07 (+) | 3.20e-07 (-) | 2.70e-01 (-) | 8.87e-09 | 1.65e-02 |
| rs6059932 | 20 | 33.2 | PIGU(0) | 1.30e-01 (+) | 5.73e-07 (+) | 6.33e-05 (-) | 6.33e-02 (+) | 1.27e-09 | 4.76e-01 |
| **HIPO-D2:** |  |  |  |  |  |  |  |  |  |
| rs4683438 | 3 | 142.7 | LOC100507389(0) | 5.70e-05 (-) | 6.85e-01 (+) | 3.77e-04 (-) | 2.87e-07 (-) | 5.41e-01 | 2.99e-08 |
| rs176813 | 4 | 69.6 | UGT2B15(+63.04kb) | 2.62e-05 (+) | 1.75e-01 (+) | 4.23e-04 (+) | 5.68e-08 (+) | 5.62e-01 | 8.10e-09 |
| rs2268719 | 6 | 52.4 | TRAM2(0) | 7.52e-07 (-) | 3.08e-01 (+) | 4.14e-02 (-) | 6.66e-08 (-) | 9.39e-01 | 2.13e-08 |
| rs7939352 | 11 | 78.0 | GAB2(0) | 2.79e-06 (-) | 9.75e-01 (-) | 5.47e-03 (-) | 2.48e-07 (-) | 9.57e-01 | 3.47e-08 |

TG: triglycerides; TC: total cholesterol. HIPO-D1 and HIPO-D2: 1<sup>st</sup> and 2<sup>nd</sup> HIPO components. The weights for the first and second HIPO components are: $\hat{\beta}_{HIPO-D1} = 0.147\hat{\beta}_{LDL} - 0.618\hat{\beta}_{HDL} + 0.591\hat{\beta}_{TG} - 0.469\hat{\beta}_{TC}$ , $\hat{\beta}_{HIPO-D2} = 0.206\hat{\beta}_{LDL} - 0.017\hat{\beta}_{HDL} + 0.228\hat{\beta}_{TG} + 0.765\hat{\beta}_{TC}$ .

### Application to psychiatric diseases

Applications of HIPO to Psychiatric Genomics Consortium (PGC) cross-disorder data^20^ show that most association signals are captured by HIPO-D1 (**Supplementary Table 6**), which has an average NCP twice larger than that of HIPO-D2. The first HIPO component puts the highest weights on BIP and SCZ, which have the largest heritability and relatively large sample sizes. It is noteworthy that for a few of the strongest signals, HIPO is outperformed by standard meta-analysis, which was implemented in PGC cross-disorder analysis as a way for detecting SNPs that may be associated with multiple traits. The QQ plot of HIPO-D1, however, dominates those for the individual traits and for the standard meta-analysis when *p* > 1×10^-8^(**Figure 1(b)**). This suggests that HIPO is superior to standard meta-analysis in detecting moderate effects, while sacrificing some efficiency for the top hits. The value of *λ*_*GC*_ and average *χ*^2^-statistics are higher for HIPO-D1 than those for individual traits and standard meta-analysis.

**Figure 1.**
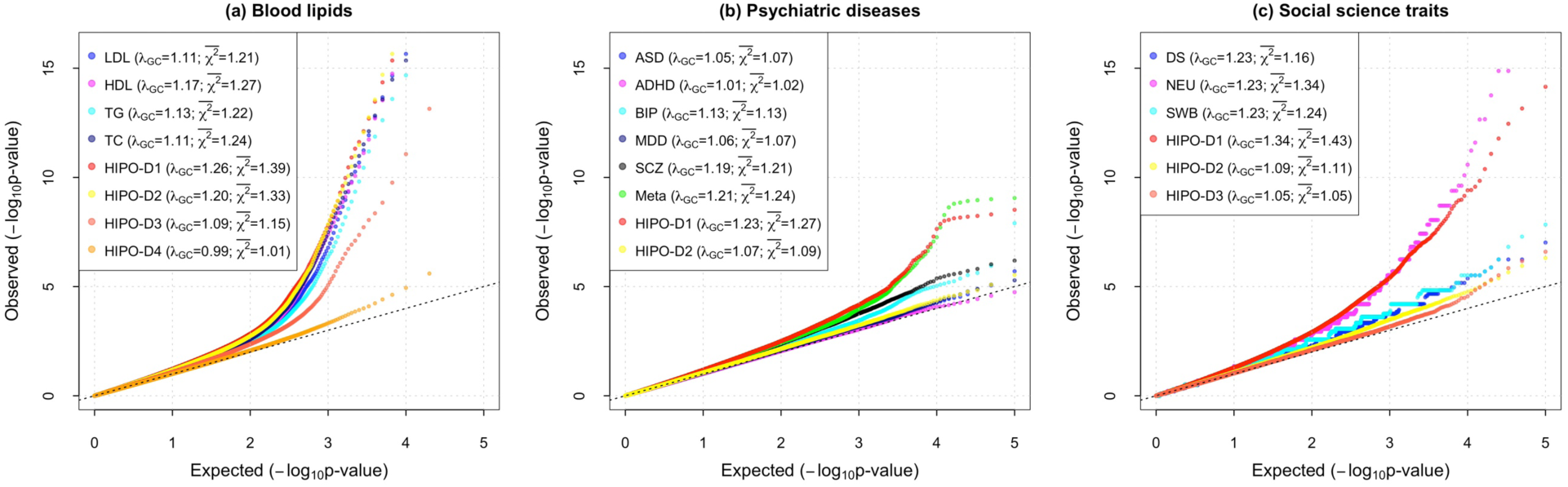
QQ plots for individual traits and underlying HIPO components across blood lipids, psychiatric diseases, social science traits. Blood lipid traits include HDL, LDL, triglycerides (TG) and total cholesterol (TC). Psychiatric diseases include autism spectrum disorder (ASD), ADHD, bipolar disorder (BIP), major depressive disorder (MDD) and schizophrenia (SCZ). Meta-analysis QQ plot is also included for psychiatric diseases (in green). Social science traits include depressive symptoms (DS), neuroticism (NEU) and subjective well-being (SWB). Genomic control factors and average *χ*^2^ statistics are shown in the legend.

HIPO-D1 discovers one new locus, marked by the lead SNP rs13072940 (*p* = 1.71× 10^-8^), that is not identified by either the individual traits or the meta-analysis. The marker rs13072940 shows association with bipolar disorder (*p*_BIP_ = 0.0026) and schizophrenia (*p*_SCZ_ = 2.55×10^-6^) but no association with autism spectrum disorder (*p*_PSD_ = 0.97), ADHD (*p* = 0.70) or major depressive disorder (*p* = 0.11). The meta-analysis signal (*p*_*M*_*eta*__ = 7.02×10^-6^) did not reach genome-wide significance and is, in fact, weaker compared to that from schizophrenia alone. This SNP show stronger association in more recent larger studies bipolar disorder^47^ (*p*_*BIP*_ = 0.0003) and schizophrenia^51^ (*p*_*SCZ*_ = 1.32×10^-7^), clearly indicating that this is likely to be a true signal underlying multiple PGC traits.

### Application to social science traits

Application of HIPO to Social Science Genetic Association Consortium studies reveals that most of the genetic variation is captured by HIPO-D1 that has an average NCP twice larger than that of HIPO-D2 (**Supplementary Table 7**). The component is negatively associated with DS and NEU and is positively associated with SWB. The tail region of QQ plot of HIPO-D1 lies close to that of neuroticism, but the values of *λ*_*GC*_ and average *χ*^2^ are substantially larger for HIPO-D1 (**Figure 1(c)**). HIPO-D1 identifies 12 new loci that are not discovered by individual trait analysis of SSGAC data (**Table 2**), increasing the total number of genome-wide significant loci from 24 to 36 (**Supplementary Figure 7**).

**Table 2.**
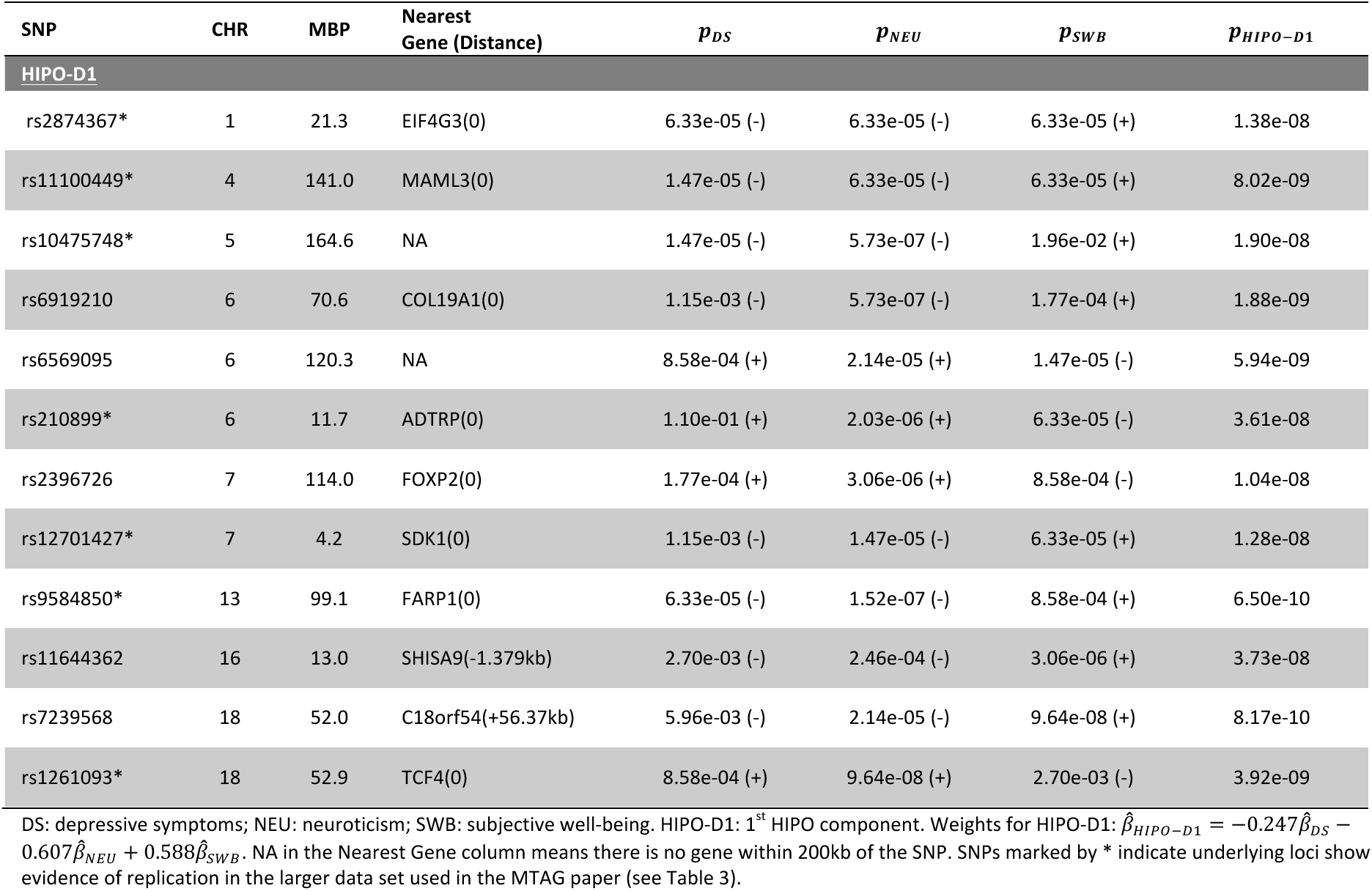
Novel loci discovered at genome-wide significance level *(p* < 5×10^-8^ by HIPO-D1 for social science traits. Independent SNPs are identified through LD-pruning with *r*^2^ threshold of 0.1 and pruned SNPs are assumed to represent independent loci if they are >0.5Mb apart. Loci are considered novel if they were not identified at genome-wide significance level through analysis individual traits. For each lead SNP, p-values for association are shown for HIPO components and for individual traits. The directions of association (+/-) are also shown for each of the individual traits.

We examined evidence of replication of the novel loci based on more recent and larger studies of DS and SWB that were incorporated in the MTAG analysis^36^. As this study reported only a list of top SNPs (*p* < 1×10^-5^) after stringent LD-pruning (*r*^2^< 0.1), we could not look up the exact lead SNPs that we report for the novel regions (**Table 2**). Instead, we searched for SNPs in the top list reported by MTAG study that could be considered proxy (D’>0.75) for our lead SNPs. We found 7 of the 12 novel have such proxies and these proxy SNPs show stronger level of association in the more recent MTAG study for at least one of DS and SWB (**Table 3**).

**Table 3.** Evidence of replication of novel loci identified by HIPO analysis for social science traits in subsequent larger studies of DS and SWB. Reported are P-values for proxy SNPs (D^’^ > 0.75) for individual trait associations in SSGAC data and the more recent MTAG study. Novel loci are identified through analysis of SSGAC which include studies of DS and SWB with sample sizes N_eff_=161,460 and N=298,420, respectively. The MTAG study includes an expanded set of sample with N_eff_=354,862 and N=388,538 for DS and SWB, respectively.

| Lead SNP in Novel Loci | Proxy SNP Reported in MTAG Study | $D'$ | Individual Trait p-value in SSGAC | Individual Trait p-value in MTAG Study |
| --- | --- | --- | --- | --- |
| **DS** |  |  |  |  |
| rs11100449 | rs1877075 | 0.78 | 2.00e-06 | 1.10e-06 |
| rs10475748 | rs10045971 | 0.99 | 4.51e-02 | 1.17e-09 |
| rs12701427 | rs4723416 | 0.91 | 1.59e-03 | 1.17e-06 |
| rs9584850 | rs4772087 | 1.00 | 2.42e-03 | 1.04e-06 |
| rs1261093 | rs11876620 | 0.82 | 1.58e-04 | 4.45e-08 |
| **SWB** |  |  |  |  |
| rs2874367 | rs12125335 | 1.00 | NA | 7.09e-08 |
| rs11100449 | rs769664 | 0.79 | 3.18e-03 | 4.59e-07 |
| rs210899 | rs10947543 | 0.93 | NA | 3.10e-08 |

DS: depressive symptoms; NEU: neuroticism; SWB: subjective well-being. NA indicates that the proxy SNP is not present in the SSGAC data.

## Discussion

In this report, we present a novel method for powerful pleiotropic analysis using summary level data across multiple traits, accounting for both heritability and sample size variations. Application of the proposed method to three groups of genetically related trait identifies a variety of novel and replicable loci that were not detectable by analysis of individual traits at comparable level of confidence. We also conduct extensive simulation studies in realistic settings of large GWAS to demonstrate the ability of the method to maintain type-I error, achieve robustness to population stratification and enhance detection of novel loci. The novel method we introduce for directly simulating summary-level GWAS statistics, preserving expected correlation structure across both traits and SNP markers, will allow rapid evaluation of alternative methods for pleiotropic analysis in settings of large complex GWAS more feasible in the future.

Application of the proposed method provides new insight into the genetic architecture of groups of related traits. For blood lipids, which have similar sample sizes, the average NCPs for HIPO-D1 and HIPO-D2 dominate the other two, suggesting that there are perhaps two unrelated mechanisms through which most genetic markers are associated with the individual cholesterol traits. For psychiatric diseases and social science traits, the top HIPO direction dominates the others, indicating that there is perhaps one major genetic mechanism underlying each group of traits. However, given that top HIPO direction down weights traits with smaller sample sizes, it is possible that there exist other independent genetic mechanisms related to these traits that could not be captured by the top HIPO component. Nevertheless, HIPO, by taking into account both heritability and sample sizes, provides a clear guideline how many independent sets of tests should be performed across the different traits to capture most of the genetic signals.

Earlier studies have proposed methods for association analysis in GWAS informed by heritability analysis. For analysis of multivariate traits observed on the same set of individuals, the MaxH^34^ method was proposed to conduct association analysis along directions that maximizes trait heritability. HIPO allows a generalization of this approach by taking into account sample size differences and overlaps across studies allowing powerful cross-disorder analysis using only summary-level data across distinct studies.

Another closely related method is MTAG^36^, which also utilizes summary level data and LD score regression to estimate genotypic and phenotypic variance-covariance matrices. MTAG, however, performs association tests for each individual trait by improving estimation of the underlying association coefficients using cross-trait variance-covariance structure. In contrast, we propose finding optimal linear combination of association coefficients across traits that will maximize the power for detecting underlying common signals. The advantage of MTAG is that it does associate the SNPs to individual traits and thus has appealing interpretation. However, strictly speaking, MTAG, similar to HIPO, is only a valid method for testing the global null hypothesis of no association of a SNP across any of the traits and may identify a SNP to be associated with a null trait while in truth it is only related to another trait in the same group. The advantage of HIPO is that it directly focuses on optimization of power in orthogonal directions for cross-disorder analysis and can provide significant dimension reduction for analysis of higher dimensional traits.

There exists a variety of methods for pleiotropic analysis^30,31,35,38,39^ that aim to optimize power for testing associations with respect to individual SNPs without informed by heritability. The method ASSET^39^, for example, searches through different subsets of traits to find the optimal subset that yields the strongest meta-analysis *z* statistic for each individual SNP. Methods like HIPO and MTAG, which use estimates of heritability based on genome-wide set of markers, are likely to be more powerful when the underlying traits have strong genetic correlation, such as that observed for psychiatric disorders. In contrast, methods such as ASSET may be more powerful for analysis of groups of traits that have more moderate genetic correlation, such as cancers of different sites^13^, for detection of loci with unique but insightful pleiotropic patterns of association. There is potential to develop intermediate methods, which borrows information across SNPs but in a more localized manner, for example, based on functional annotation information^52,53^.

In conclusion, HIPO provides a novel and powerful method for joint association analysis across multiple traits using summary-level statistics. Application of the method to multiple datasets shows that it provides unique insight into genetic architecture of groups of related traits and can identify substantial number of novel loci compared to analysis of individual traits. Further extension of the method is merited for facilitating more interpretable and parsimonious association analysis across groups of high-dimensional correlated traits.

